# Araport11: a complete reannotation of the *Arabidopsis thaliana* reference genome

**DOI:** 10.1101/047308

**Authors:** Chia-Yi Cheng, Vivek Krishnakumar, Agnes Chan, Seth Schobel, Christopher D. Town

**Author notes:** These authors contributed equally to this work. Corresponding author (C.D.T.).

## Abstract

The flowering plant *Arabidopsis thaliana* is a dicot model organism for research in many aspects of plant biology. A comprehensive annotation of its genome paves the way for understanding the functions and activities of all types of transcripts, including mRNA, noncoding RNA, and small RNA. The most recent annotation update (TAIR10) released more than five years ago had a profound impact on *Arabidopsis* research. Maintaining the accuracy of the annotation continues to be a prerequisite for future progress. Using an integrative annotation pipeline, we assembled tissue-specific RNA-seq libraries from 113 datasets and constructed 48,359 transcript models of protein-coding genes in eleven tissues. In addition, we annotated various classes of noncoding RNA including small RNA, long intergenic RNA, small nucleolar RNA, natural antisense transcript, small nuclear RNA, and microRNA using published datasets and in-house analytic results. Altogether, we identified 738 novel protein-coding genes, 508 novel transcribed regions, 5051 non-coding genes, and 35846 small-RNA loci that formerly eluded annotation. Analysis on the splicing events and RNA-seq based expression profile revealed the landscapes of gene structures, untranslated regions, and splicing activities to be more intricate than previously appreciated. Furthermore, we present 692 uniformly expressed housekeeping genes, 43% of whose human orthologs are also housekeeping genes. This updated Arabidopsis genome annotation with a substantially increased resolution of gene models will not only further our understanding of the biological processes of this plant model but also of other species.

## INTRODUCTION

Adopted by the research community over 50 years ago as a model for plant research (Rédei 1975; Provart et al. 2016), *Arabidopsis thaliana*, a member of the crucifer family, continues to occupy a prominent place in plant biology. It also has an underappreciated influence on medical research and human health. Studies using *Arabidopsis* have played a leading role in basic biological discoveries (Jones et al. 2008) such as the plant NB-LRR proteins and their later identified human orthologs in the innate immune system (Jones and Dangl 2006), the impact of auxin research on the ubiquitin pathway conserved among eukaryotes (Parry and Estelle 2006), a light signaling component COP1 whose mammalian orthologs has a role in tumorigenesis (Deng et al. 1991), and more.

The *Arabidopsis thaliana* genome sequence was initially assembled and annotated in 2000 by an international consortium (Arabidopsis Genome Initiative 2000). The sequence derived from the Columbia-0 (Col-0) ecotype was subsequently revised twice culminating in the TAIR9 Genome. Refined annotation sets were released by The Institute for Genomic Research (TIGR, versions 1-4) (Haas et al. 2005) and subsequently by The Arabidopsis Information Resource (TAIR, versions 5-10) (Lamesch et al. 2012). The TAIR10 annotation was informed by *ab initio* gene models, EST sequences from Sanger platforms, and two RNA-seq datasets available at that time. Since TAIR10, nearly two hundred *Arabidopsis thaliana* RNA-seq studies have been published and deposited in NCBI SRA. In comparison to the EST data that provided the bulk of the TAIR10 annotation, the RNA-seq data offer single-base resolution and more precise measurement of levels of transcripts and their isoforms (Wang et al. 2009). Thus, the publically available RNA-seq datasets present a compelling opportunity to update the TAIR10 annotation. The literature since TAIR10 reveals a growing amount of information about noncoding RNA, including long intergenic RNA, natural antisense transcript, small RNA, microRNA, small nuclear RNA, small nucleolar RNA and tRNA (Sherstnev et al. 2012; Wang and Brendel 2004; Kozomara and Griffiths-Jones 2014; Okamoto et al. 2010; Matsui et al. 2008; Csorba et al. 2014; Li et al. 2013; Liu et al. 2012). As we begin to understand their regulatory roles in growth and development, a complete documentation and uniform annotation of noncoding RNAs will serve as a platform for the further progress.

We report an update to the annotation of the *Arabidopsis thaliana* Col-0 genome. The new annotation offers refined gene structures applied to the TAIR10 genome sequence and updated functional descriptions for many genes. In an effort to exclude false gene models formed by combining isoforms, the analysis is based on tissue-specific assemblies of RNA-seq data. The new annotation is called Araport11 to maintain version number continuity and to designate the source as Araport, an open-access online resource for Arabidopsis research community (Krishnakumar et al. 2015). Araport11 will be accessible from the Araport portal (www.araport.org) and NCBI GenBank.

## RESULTS

### Deep transcriptome sequencing reveals a refined annotation with diverse transcript structure

#### Protein-coding RNA

We interrogated 113 RNA-seq datasets generated from untreated or mock-treated wild-type Col-0 plants. To comprehensively capture the transcriptome throughout the plant developmental cycle, we partitioned the RNA-seq datasets into 11 groups according to their tissue or organ of origin. These include the aerial part, carpel, dark-grown seedling, leaf, light-grown seedling, pollen, receptacle, root, root apical meristem, stage 12 inflorescence, and shoot apical meristem. We will refer to these sample sources as tissues hereafter following a previous convention (Schmid et al. 2005). Reads from each library were aligned to the TAIR10 genome assembly as described in Methods. The mapping generated 66.1 Gb of uniquely mapped sequence representing 491-fold coverage of the TAIR10 genome. These data provided 2182fold coverage of the Araport11 transcriptome (Supplemental Table S1). We used a pipeline that combined *de novo* and genome-guided Trinity assemblies (Grabherr et al. 2011) followed by PASA (Haas et al. 2003) to assemble maximal length transcripts. In order to instantiate tissue-specific transcript isoforms and avoid chimeras between tissues, we assembled each tissue collection individually. Eleven tissue-specific assemblies were independently used to refine the augmented TAIR10 annotation (See Supplemental Methods), and we obtained a final annotation of 27,688 protein-coding loci with 48,359 transcripts (Table 1A).

**Table 1.** Summary of Araport11

| Type | TAIR10 | Araport11 |
| --- | --- | --- |
| (A) Protein-coding gene |  |  |
| Number of loci | 27,416 | 27,688* |
| Number of transcripts | 35,386 | 48,359 |
| Number of loci with two or more splice variants | 5,804 | 10,696 |
| (B) Noncoding gene |  |  |
| Long intergenic noncoding RNA (lincRNA) | 36 | 2,495 |
| Natural antisense transcript (NAT) | 223 | 1,115 |
| microRNA (miRNA) | 177 | 325<sup>Γ</sup> |
| Small nucleolar RNA (snoRNA) | 71 | 287 |
| tRNA | 689 | 714<sup>Ψ</sup> |
| Small nuclear RNA (snRNA) | 13 | 82 |
| rRNA | 15 | 15 |
| Unclassified | 394 | 18 |
| (C) Genomic feature |  |  |
| Small RNA |  | 35,846 |
| Novel transcribed region |  | 508 |
| Upstream open reading frame | 58 | 84 |
| Obsolete loci with short coding sequence |  | 388 |

\* A total of 747 novel loci and 388 obsoleted loci and some gene splits and merges (see Supplemental Methods for details).
<sup>Γ</sup> A total of 427 mature miRNAs derived from 325 primary transcripts
<sup>Ψ</sup> This includes 689 tRNA loci and 25 pseudogenic loci.

To update the functional annotation of novel and existing protein-coding loci, we used a weighted keyword (WK) approach. In brief, each predicted protein sequence was searched against databases (Priam, Uniref100, PFAM/TIGRFAM, CAZY, CDD) and scanned with motif finders (TMHMM, InterPro). Keywords were extracted from the definition lines of best matches and scored based on a set of heuristic rules. The maximal scoring definition line became the functional annotation if the score surpassed a threshold. This process generated functional annotations of 7,122 protein-coding loci including 747 loci not annotated in TAIR10.

#### Noncoding RNA

We incorporated the curated datasets from miRBASE (Kozomara and Griffiths-Jones 2014) and PlantRNA database (Cognat et al. 2013) as well as characterized genes (e.g. *COOLAIR*) (Csorba et al. 2014) in Araport11. For the long-intergenic RNAs, we obtained the published results (Liu et al. 2012) and annotated 2,704 loci that fall into the intergenic regions in Araport11. The accurate detection of natural antisense transcripts relies upon strand-specific sequencing technology combined with statistical-computational approaches. It has been suggested that depending on the protocol used for strand-specific RNA library preparation, 0.5% to 11.2% of the reads encoded from the sense strands could be mistakenly mapped to the antisense strand (Levin et al. 2010). Therefore, we annotated the 1,115 NAT loci identified by combined strand-specific sequencing data and statistical analyses (Li et al. 2013). Because Li *et al* did not propose transcript structures, we manually curated those NAT loci using PASA-assembled transcripts. Overall, Araport11 represents a total of 5,051 noncoding RNA loci (Table 1B).

#### Pseudogene

The mapped RNA-seq data indicated that many pseudogenic loci are transcribed: among the 924 loci annotated as pseudogene in TAIR10, 259 were supported by our transcript assemblies. We manually curated 425 pseudogenic transcript models on these loci using Web Apollo (Lee et al. 2013). Over 30% of the transcribed pseudogenic loci encode two or more transcript variants, many of which have clearly defined spliced structures (Supplemental Fig. S1). The level of splicing activity is comparable to that of protein-coding genes (38%), which is intriguingly complex for genomic features traditionally considered non-functional. Furthermore, we updated the functional annotation for 882 pseudogenes by using blastx to find the best matched counterparts in Arabidopsis proteins. We consolidated this dataset with 25 pseudogenic tRNAs curated by the PlantRNA database (Cognat et al. 2013) and obtained a final annotation of 952 pseudogenes in Araport11.

#### Upstream open reading frame

Upstream open reading frames (uORFs) are common genomic features in the 5’ untranslated regions of eukaryotic mRNAs. In Arabidopsis, identification of uORFs has previously relied on the conserved peptides encoded by the uORFs (CPuORFs). These cases are thought to represent less than 1% of all uORFs in plants and animals. In total, 64 CPuORFs associated with 58 loci (Hayden and Jorgensen 2007) were annotated in TAIR10, while thousands more uORFs have been implicated by computational prediction and ribosome profiling data (Takahashi et al. 2012; Juntawong et al. 2014). Araport11 includes additional 26 uORFs from the literature, each with characterized biological functions (Laing et al. 2015; Saul et al. 2009; Takahashi et al. 2012; Rosado et al. 2012) (Table 1C). Noting that one main ORF may have more than one uORF, we adopted a nomenclature (e.g. AT1G67480.uORF1) that associated the uORF with its cognate protein-coding locus and assigned an isoform number. Additional 6,680 loci with predicted uORFs bound by ribosomes at a significant level and have yet to be annotated are publicly available via JBrowse tracks at Araport (Bailey-Serres, Bazin, and Girke, personal communication).

### Evaluation of annotation

A quantitative evaluation of the accuracy of the exon-intron structure is a critical step toward maintaining a gold standard annotation. To this end, we used annotation edit distance (AED) (Eilbeck et al. 2009) to assess Araport11 annotation. AED is generated by an actively maintained software MAKER-P and has been shown to closely correspond to the confidence classification used by TAIR (Campbell et al. 2014). AED measures the consistency of gene models with the available nucleotide and protein sequence alignments. Each transcript was assigned a computed AED score, between 0 and 1, with 0 denoting complete agreement with the evidence and 1 indicating complete absence of supporting evidence. A comparison of the AED score distribution showed that overall Araport11 gene models are in closer concordance with the underlying evidence than TAIR10 models (Fig. 1A).

**Figure 1:**
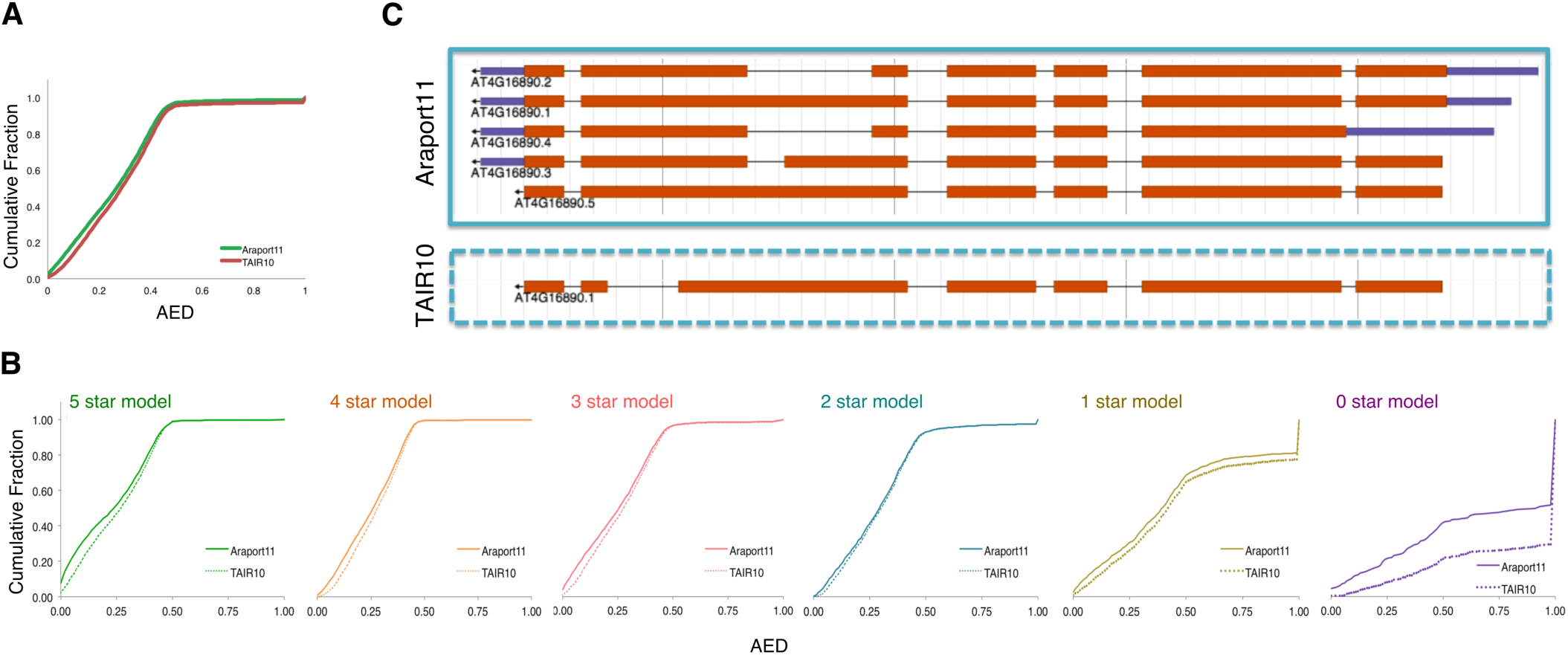
Accuracy of annotation. Annotation Edit Distance (AED) measures the consistency of transcript models with the underlying nucleotide/protein alignments. Lower AED scores suggest that the structures are in better agreement with the evidence. The cumulative fraction of AED scores provides a quantitative means to evaluate the annotation of all genes (A) which were broken down into 0 to 5 star ranks (B) according to TAIR10 annotation. (C) An example of gene structure improvement made to AT4G16890.1, which was previously annotated with an erroneous 5th intron. In addition to the gene structure correction, we also added novel isoforms assembled via the Araport11 pipeline.

We examined the cumulative fraction of AED in each category of the TAIR10 five-star ranking system (Fig. 1B). For the lowly ranked genes, one obvious improvement was made for the zero star models, which, at the time of the TAIR10 annotation, did not have supporting evidence. Many updates were made to the TAIR10 transcripts that had been classified as two stars or one star, signifying that the previously available evidence did not completely cover the junctions or coding regions. One example, AT4G16890.1, is a two-star model previously annotated with an erroneous 5th intron. In Araport11, we corrected the gene structure (Fig. 1C) and added novel isoforms detected in multiple tissues. Importantly, the improvements made in Araport11 were not limited to low-confidence models but rather across all annotation classes. This is because the highly ranked models had more expression evidence available as raw materials for refinement. As another example, AT2G36480.1 and AT2G36485.1, two five star transcripts, were merged into a single transcript for the reads supporting the linking junction were present in multiple tissues.

### Small RNA

Small RNAs cover nearly 10% of the Arabidopsis genome yet have been disproportionately annotated (Coruh et al. 2014). Most efforts have focused on miRNAs that constitute a minor portion of the small RNA repertoire, resulting in a gap between the knowledge of small RNA expression and the annotation of small RNAs in Arabidopsis (Coruh et al. 2014). We analyzed small RNA-seq datasets generated from root (Hsieh et al. 2009; Breakfield et al. 2012), leaf (Yu et al. 2013), aerial part (Fahlgren et al. 2010), flower (Lister et al. 2008; Cuperus et al. 2010; Law et al. 2013), embryo (Lu et al. 2012), and silique (Hardcastle et al. 2012) (Supplemental Table S2). We identified *de novo* small RNA clusters with ShortStack (Axtell 2013) that has been used to annotate small RNA genes in plant and animal species (Jex et al. 2014; Lunardon et al. 2016; Coruh et al. 2015). This method yielded a total of 35,846 clusters which predominantly produced 24-nt small RNAs (Fig. 2A). Consistent with previous findings (Zhang et al. 2007; Law et al. 2013), most of these small RNA clusters are PolIV-dependent (Fig. 2B). They also tend to overlap with transposable element-related regions (Fig. 2C). We compared our results with published genomic regions that generate PolIV-dependent RNAs (P4RNAs), the precursors of small RNAs (Zhai et al. 2015; Li et al. 2015). Our clusters completely recapitulated the 7,631 200-bp static regions that generated small RNAs in stage-12 inflorescence (Law et al. 2013), later shown to encode P4RNAs (Zhai et al. 2015). Furthermore, the ShortStack-identified clusters overlapped with 95% of the P4RNA loci (Fig. 2D) independently identified by another group using unopened flowers (Li et al. 2015). We hypothesized that the diverse tissue types used in our pipeline contributed to the clusters exclusively detected in this work. Indeed, ~43% of the small RNA clusters were detected only in non-flower tissues. In summary, we analyzed more than 124 million mapped small RNA-seq reads and constructed a comprehensive set of small RNA generating loci. The small RNA annotations, including the genomic location and underlying metadata, can be interactively explored via JBrowse at Araport and are also available in Supplemental Data Set S1.

**Figure 2:**
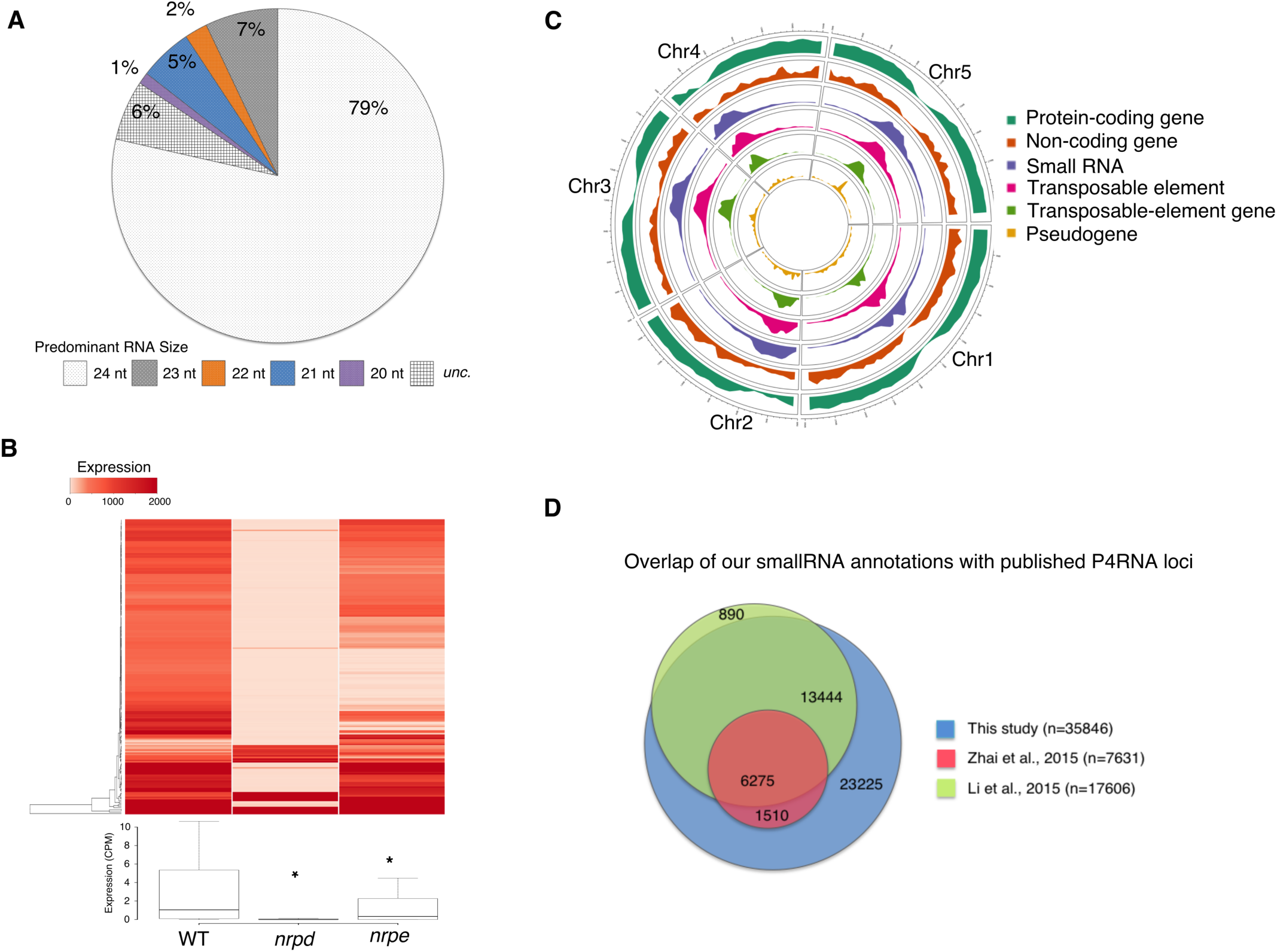
Properties of small RNAs. (A) Size distributions within 35,846 small RNA loci. *unc*., uncharacterized means that a majority of RNA size is lacking for those loci. (B) Heatmap and boxplot of small RNA levels comparing the wild-type, *nrpd* (*pol-iv*) and *nrpe* (*pol-v*) flowers (* indicates significant reduction; *P*-value < 2.2 × 10^−16^, Wilcoxon rank sum test). (C) Circular representation of genome-wide distribution of genomic features as indicated by the legend. (D) A Venn diagram showing the overlap of small RNA generating loci (this study) and P4RNA loci (Li et al. 2015; Zhai et al. 2015).

### Diversity of alternative splicing

Nearly 40% of Araport11 protein-coding loci encode two or more alternative splicing isoforms. In *Drosophila* and mammals, over 57% and 95% of genes produce more than one transcript isoforms respectively (Wang et al. 2008; Pan et al. 2008; Brown et al. 2014). Given the depth of our RNA-seq datasets, the gap likely reflects the nature of different organisms. While the majority of the protein-coding loci are transcribed into two to three different variants, there are 388 loci capable of encoding between seven to twenty-seven isoforms. These 388 genes are overrepresented in the organic cyclic compound metabolic process (GO:1901360), nucleic acid metabolic process (GO:0090304), and cellular aromatic compound metabolic process (GO:0006725), implying a correlation between versatile transcript structures and these metabolic processes. In line with previous findings (Palusa et al. 2007), the serine/arginine-rich proteins, a family of splicing regulators conserved in eukaryotes, are also overrepresented in these 388 loci with high numbers of isoforms.

To examine the general nature of alternative splicing (AS), we further characterized the AS events using the SUPPA software (Alamancos et al. 2015) and obtained a total of 19,915 splicing events in Araport11 gene models, over 65% of which were not annotated in TAIR10. The AS events consist of 10,375 retained introns (RI), 4,230 alternative 3’ splice sites, 3,172 alternative 5’ splice sites, 1,051 skipped exons, 944 alternative first exons, 113 alternative last exons, and 30 mutually exclusive exons (Fig. 3A). To examine the variation of splicing events throughout development, we computed a “percent splicing index” (PSI or ψ) across the eleven tissues. PSI is a value that denotes the efficiency of a given splicing event and calculated as the fraction of the supporting isoform(s) to the total isoforms. The distribution of PSI across tissues revealed the versatile nature of splicing activity (Fig. 3B). Unsupervised hierarchical clustering revealed that many splicing events are specific to particular tissue type(s) and likely to be regulated in a tissue-specific manner.

**Figure 3:**
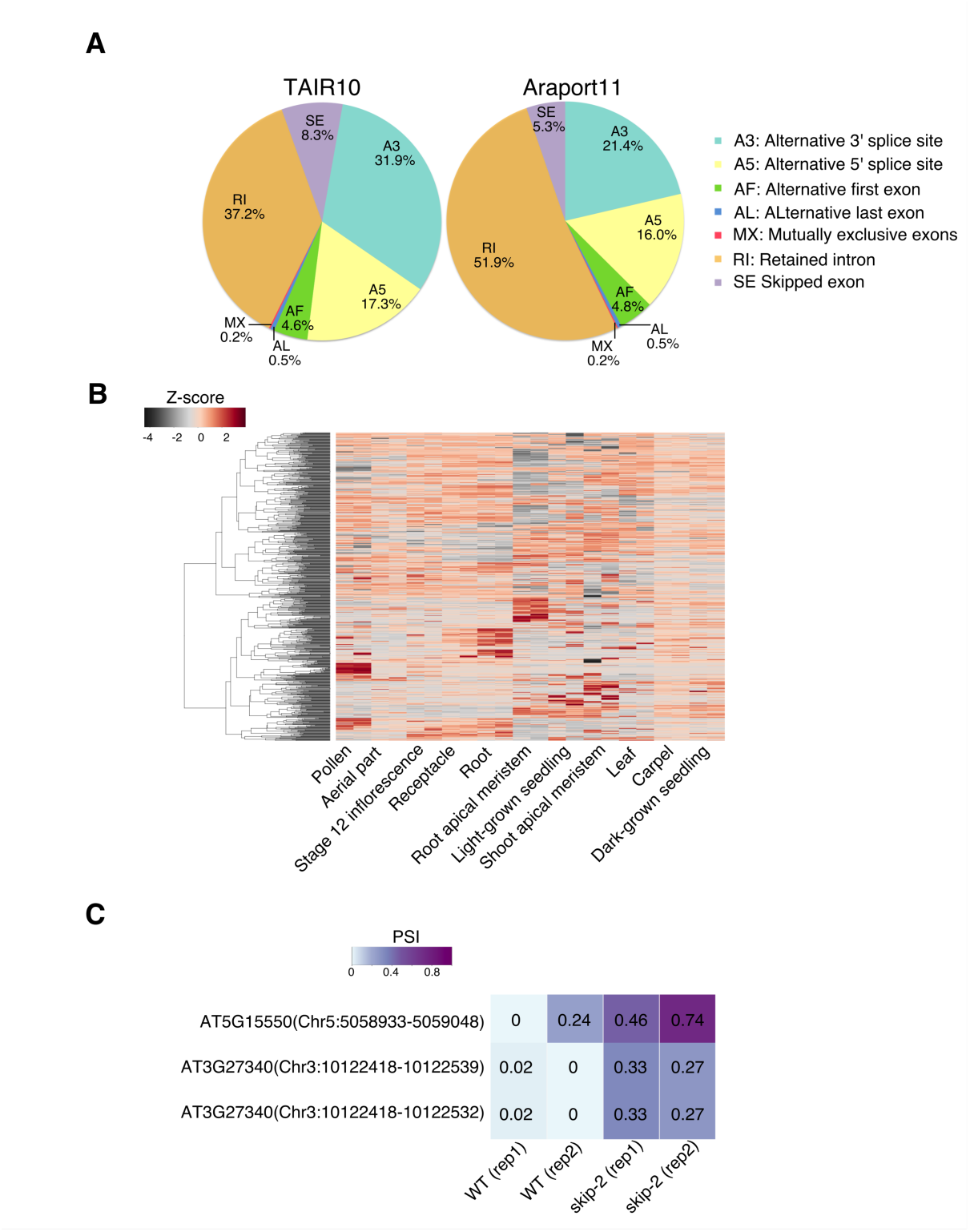
Features of splicing events. (A) Pie charts showing proportions of different classes of alternative splicing events in the Araport11 and TAIR10 annotations. (B) Unsupervised clustering of splicing events across eleven tissues. The scale bar indicates z-scores of ψ. (C) Comparison of ψ for three experimentally verified retained introns between wild-type and splicing-defective *skip-2* plants.

Studies using various experimental approaches and computational analyses have conclusively indicated that intron retention is the most prevalent AS event in Arabidopsis and rice, ranging from 40% to 65% (Ner-Gaon et al. 2004; Iida et al. 2004). This is different from human AS events, in which intron retention represents the least common (3.5%) type of event. We explored the dependency of retained introns on spliceosome activity by calculating the PSI of RI events in wild type and a *skip-2* mutant defective in a splicing factor conserved in plants, yeast, and human. We computed the PSI values of the retained introns whose abundance increased in *skip-2* as verified by RT-PCR (Wang et al. 2012) and found that their PSI values also increased in *skip-2*. This demonstrated the concordance of PSI values with experimental evidence of transcript abundance (Fig. 3C). We extended the analysis to a genome-wide level and found that compared with wild type, the PSI values for 6,641 RI events in *skip-2* were significantly higher (Wilcoxon rank sum test, *P*-value < 2.2 x 10^−16^), suggesting a positive role of spliceosomes in the recognition and removal of these introns.

Nonsense-mediated decay (NMD) is a conserved eukaryotic quality-control mechanism that eliminates both normal and aberrant transcripts with premature termination codons (Shaul 2015). Many intron-retained transcripts were not sensitive to NMD as their transcript abundance did not increase in a *upf1 upf3* mutant defective in NMD (Kalyna et al. 2012). We systematically examined the relations of NMD and intron retention by comparing the PSI values for wild-type and *upf1 upf3* seedlings using published RNA-seq datasets (Drechsel et al. 2013). Our analysis showed that the overall PSI values for 5,499 RI events were reduced in *upf1 upf3* (Wilcoxon rank sum test, *P*-value = 0.0005112), suggesting that the intron-retained transcripts were not necessarily targeted by NMD in Arabidopsis.

### The exitrons are occupied by ribosomes

About 17% (1,504) of the retained introns in Araport11 are exonic introns, or exitrons, which are introns where “both splice sites inside an annotated coding exon” (Marquez et al 2015). This subset of introns tends to be shorter and flanked by weaker splice sites in vertebrates and plants (Zavolan et al. 2003; Berget 1995). The impact of exitrons on the coding sequence of the transcript is dependent on their lengths (Fig. 4A-D). Nearly 60% of the exitrons have the lengths divisible by three (EI_x3_); therefore, retaining EI_x3_ does not alter the reading frame and will only increase the length of the coding sequence. For exitrons with lengths not divisible by three (non-EI_x3_), retention does not necessarily introduce a stop codon downstream from the splice junctions (Fig. 4C-D). Taken together, our analyses show that over 80% of the exitron-retained transcripts have longer coding sequences than their exitron-spliced counterparts. We then compared the ribosome occupancies of exitrons, introns and coding exons within the same locus. The ribosome occupancy was calculated as the ratio of the relative abundance of each feature (intron, exitron, or coding exon) in the ribosome footprint library (Liu et al. 2013) to those of the control RNA library. We found that in etiolated seedlings, the ribosome occupancies of exitrons were significantly higher than those of introns (Fig. 4E).

**Figure 4:**
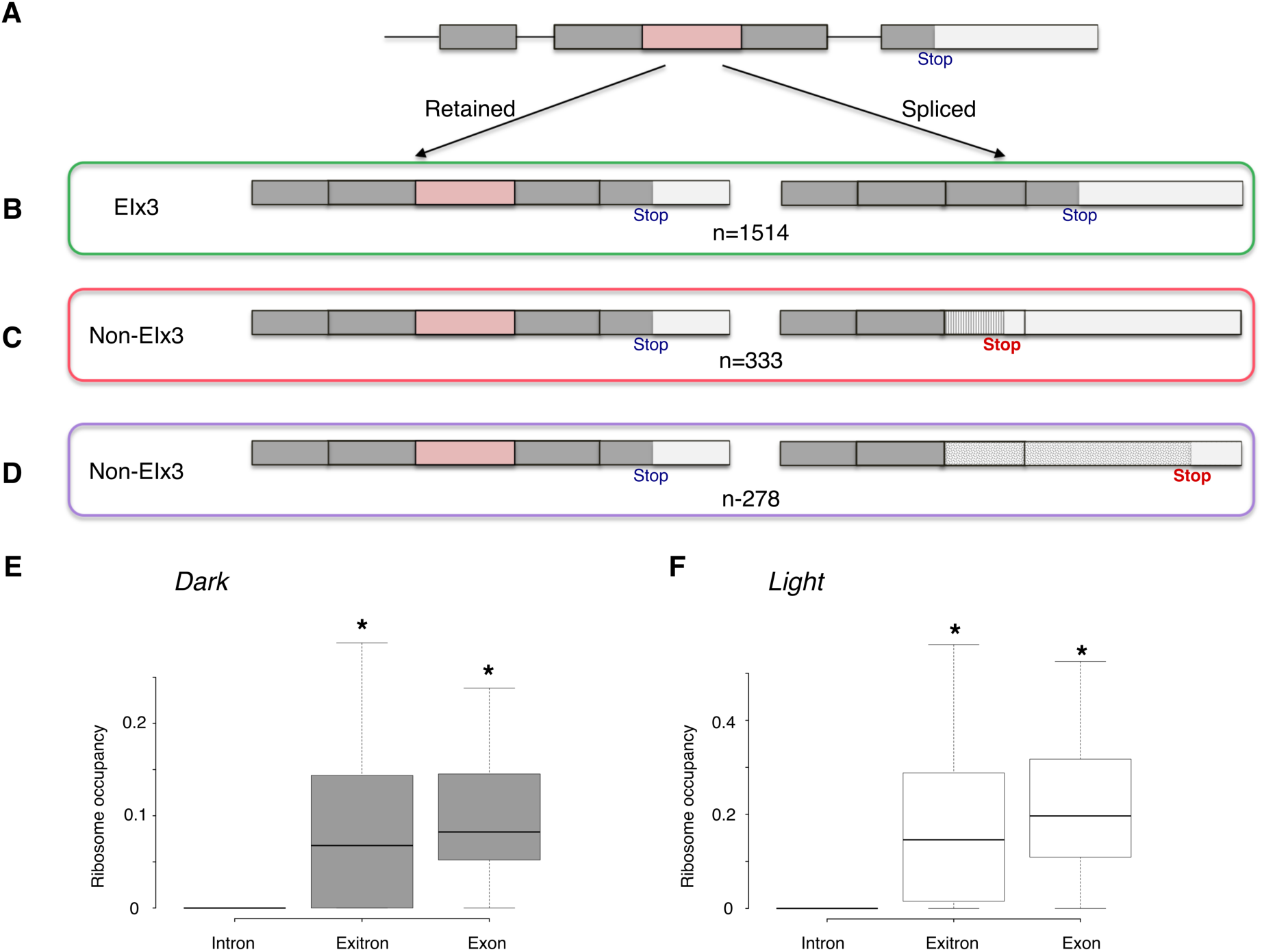
Exitrons. (A) Both splice sites of an exitron (pink) are localized within the coding region (shaded rectangles). The impact of exitron retention or splicing on the CDS is dependent on the length of the exitron (B-D). (B) When the exitron length is a multiple of three, splicing out an exitron results in the deletion of coding sequence. When the exitron length is not a multiple of three, splicing out an exitron changes the C-terminal coding sequence and the new stop codon could be upstream (C) or downstream (D) in comparison to the exitron-retained transcript. (E,F) Boxplots showing the ribosome occupancies for etiolated seedling without (E) and with (F) exposure to light (* indicates significant increase; *P*-value < 2.2 × 10^−16^, Wilcoxon rank sum test).

### Discovery of novel transcribed regions

Incorporation of predicted protein-coding genes from the Gnomon (Souvorov et al. 2010) and MAKER-P (Campbell et al. 2014) pipelines resulted in the instantiation of 738 novel protein-coding genes. Over 96% of these novel genes were supported by our RNA-seq data. We further identified 508 intergenic regions of Araport11 to which RNA-seq data could be mapped and designated these novel transcribed regions (NTRs). Over 60% of the NTRs encode multi-exonic transcripts, some of which are transcribed into two or more isoforms with splice variants (Fig. 5A). Furthermore, over a quarter of the NTRs contained a transcription start site reported in a large-scale analysis using whole root samples (Morton et al. 2014), providing additional support for their structures. Most novel transcripts were expressed at low levels and/or only in a limited number of tissues (Fig. 5B), which may explain why they escaped prior annotation.

**Figure 5:**
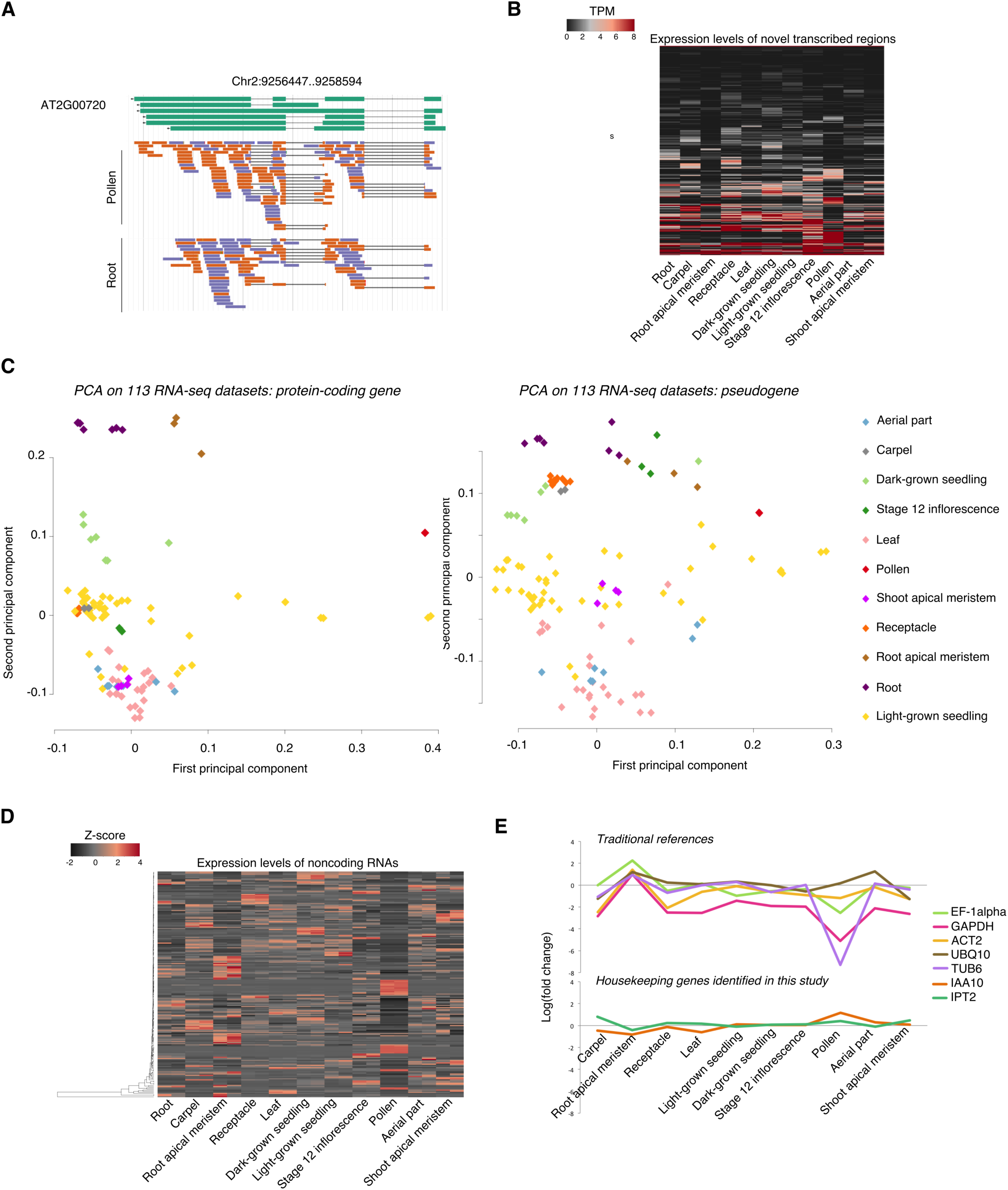
Genome-wide expression profile of several classes of transcripts. (A) An example of novel transcribed region that encodes multiple splice variants. (B) Heat map showing the expression levels of 508 novel transcribed regions across eleven tissues. (C) PCA of protein-coding genes and pseudogenes on 113 RNA-seq datasets revealed clusters sharing similar expression signatures. Note the loose cluster formed by light-grown seedling samples (yellow). (D) Heat map showing the expression of noncoding RNAs across eleven tissues. The scale bar indicates z-score of TPM. (E) Expression levels of newly identified (IAA10, IPT2) or conventionally used (EF-1α, GAPDH, ACT2, UBQ10, TUB6) housekeeping genes. Fold change shown in log_2_ scale.

Some novel transcripts have sequence similarity to known plant proteins. However, less than a quarter of these transcripts cover over 70% of the subject proteins, most of which are annotated as uncharacterized. It is plausible that these NTRs that partially cover the subject proteins could encode small peptides, be pseudogenes or noncoding RNAs. To further explore these possibilities, we examined the ribosome occupancies of the NTRs using ribosome footprinting datasets generated from light-grown and etiolated seedlings (Juntawong et al. 2014; Liu et al. 2013). In 281 NTRs we observed aligned reads associated with ribosomes, implying a likelihood of translation. In addition, we also compared the NTRs with an atlas of about 90,000 conserved noncoding sequences (CNS) generated from nine-way genome alignment of Brassicaceae species with *Arabidopsis thaliana* (Haudry et al. 2013) and found 123 NTRs located in those conserved noncoding regions. Taken together, our initial analyses have revealed the structures and expression profiles of these novel transcribed regions. We have assigned these regions unique AGI identifiers without specifying their protein-coding or noncoding classification as the diagnostic evidence is lacking. Whether they function as (small) peptide-coding RNAs or noncoding RNAs remains an open question for the future.

### Expression dynamics of tissue-specific transcriptomes

Over 99% of all genes, including 27,596 protein-coding genes, 944 pseudogenes, and 4,868 noncoding genes were detected in at least one of the 113 RNA-seq datasets, demonstrating that the use of extensive RNA-seq data improved the overall sensitivity of detection in comparison with microarrays (Schmid et al. 2005). The less than 1% of genes without detectable expression include 103 noncoding RNAs, 44 hypothetical proteins, 16 pseudogenes, and a few published/curated proteins. Inspection of additional RNA-seq datasets showed that a subset of these genes was detectable under various experimental regimes such as hormone treatment, pathogen infection, and/or nutrient stress, suggesting the expression of these genes is specific to environmental stimuli.

To comprehensively explore the gene expression dynamics throughout development, we separately examined the expression profiles of protein-coding genes, noncoding genes, and pseudogenes. We will refer to expressed genes as those with greater than one transcript per million (TPM) for the remaining analyses and describe the expression features of these three categories of transcripts in the following sections.

#### Protein-coding gene

Over 84% of protein-coding genes were expressed in one or more tissue. About 59% to 70% of the total genes were expressed in most tissues except for pollen, in which only 23% of genes were expressed. Furthermore, nearly 20% of the expressed genes are found in all eleven tissues and this rises to 57% when pollen is excluded, indicating a significant overlap of genes shared by morphologically distinct tissues. Despite this sharing phenomenon, each tissue maintains a transcriptional fingerprint as the anatomical similarity was reflected in principal component analysis (PCA). Leaf samples, for example, clustered together with each other and with samples made from the aerial part and shoot apical meristem, regardless of the growth conditions or harvesting age. Notably, light-grown seedling samples, the most widely used materials, appeared to be the most heterogeneous despite the similar plate-grown conditions and harvesting age (Fig. 5C).

Next, we explored the tissue-specific genes that were expressed in only one tissue. Pollen has the highest percentage of tissue-specific genes despite the smallest size of transcriptome; the reproductive tissues and root have higher fractions of tissue-specific genes (Supplemental Table S3). We performed GO enrichment analyses on each set of tissue-specific genes and found enriched GO terms in several tissues. For example, in the receptacle of stage 15 flowers where abscission zone cells responsible for programmed separation of plant organs were dominant, we found overrepresented GO terms related to cell death and response to stimulus in the tissue-specific genes. A complete list of tissue-specific genes and the enriched GO terms is shown in Supplemental Data Set S2.

#### Pseudogene

The sensitivity and depth of RNA sequencing is a primary factor affecting the detection of the mostly lowly expressed pseudogenes (Kalyana-Sundaram et al. 2012). Our analysis detected expression (TPM>0) of over 98% of the pseudogenes, an increase from 2%-36% using EST, massively parallel signature sequencing (MPSS), and microarray evidence (Zou et al. 2009).

Most pseudogenes are, however, lowly expressed as only 27% of pseudogenes are expressed at the level of TPM above 1 in one or more tissues. The transcription of pseudogenes also maintained a tissue-specific signature as shown by PCA analysis (Fig. 5C).

#### Noncoding gene

While 4,868 noncoding genes were detected at levels above zero TPM, only 1,644 reached a level above one TPM. On average, each noncoding gene was expressed in 5 tissues, compared to 8 tissues for protein-coding genes. Furthermore, many noncoding genes appeared to have peak expression in specific tissues (Fig. 5D); nearly four hundred were expressed in only one tissue, predominantly reproductive tissues and root. The preferential expression of noncoding genes in reproductive organs was also observed in *Drosophila* (Brown et al. 2014). Recent studies have provided some insight into the responsiveness of long intergenic RNA and natural antisense RNA to environmental stimuli (Liu et al. 2012; Wang et al. 2014). We analyzed the expression profiles of noncoding genes annotated in Araport11 in response to abscisic acid (ABA) treated seedlings (GSE65016) and identified 101 noncoding genes that are differentially expressed in 12-day-old seedling after ABA treatment. We further examined the expression profile of these ABA-regulated noncoding genes in root and shoot after mild salt treatment (Sani et al. 2013) and found groups of noncoding RNAs that are co-expressed or co-regulated in a tissue-specific manner (Supplemental Fig. S2). AT4G09695, for example, is specifically up-regulated by salt priming in the root sample.

### Identification of housekeeping genes

Uniformly expressed genes are crucial internal references for large expression datasets, and can also shed light on the mechanisms underlying basic cell maintenance. To identify the genes that are stably expressed in all tissues, we used two RNA-seq datasets per tissue type because the numbers of RNA-seq datasets for each tissue ranged from 2 to 45, thus avoiding any analysis bias brought about by favoring the more highly represented tissues. We used RUVseq (Peixoto et al. 2015; Risso et al. 2014) to remove technical effects to accommodate the heterogenous sources of the RNA-seq datasets. The adjustment of unwanted variation, including batch effect, library preparation, and other nuisance effects was followed by an ANOVA-like test in edgeR (McCarthy et al. 2012) to identify genes whose expression levels were not significantly different among any of the tissues. We further restricted the variations between tissues to be less than three fold. Overall, we identified 692 housekeeping genes consisting of 666 protein-coding genes, 12 noncoding RNAs, 6 pseudogenes, 6 transposable element genes, and 2 novel transcribed regions (Supplemental Data Set S3).

This group of housekeeping genes was enriched in pentatricopeptide repeat (IPR002885) and tetratricopeptide-like helical domain (IPR011990). GO terms associated with basal biological processes such as single-organism intracellular transport, establishment of localization in cell, vesicle-mediated transport, organelle organization, and cytoplasmic transport were also overrepresented. Additional housekeeping genes include elements involved in auxin signaling, auxin transport, and cytokinin biosynthesis, consistent with the essential roles of these two phytohormones.

The 692 housekeeping genes include 12 of the previously reported top 100 most stably expressed genes (Czechowski et al. 2005). The marginal overlap may be due to the differences in the source data (RNA-seq vs. microarray) and the identification criteria. Furthermore, the traditional reference genes, *ACT2* (AT3G18780), *TUB6* (AT5G12250), *EF-1α* (AT1G07920), *UBQ10* (AT4G05320), and *GAPDH* (AT3G04120) fluctuate across different tissues (Fig. 5F) and are not present in our list. It is noteworthy that 297 of these 692 housekeeping genes have human orthologs also characterized as housekeeping genes with uniform expression across 16 human tissues in RNA-seq data (Eisenberg and Levanon 2013), indicating a considerable conservation of core processes between plant and mammalian systems.

## DISCUSSION

Continuous refinement and routine updates of annotation are prerequisites for correctly interpreting the functional elements of the genome. Even for well-annotated genomes such as Arabidopsis, fly, human, mouse, rice, and yeast, the advantages of RNA-seq have offered an unprecedented resolution of their transcriptomes (Nagalakshmi et al. 2008; Wilhelm et al. 2008; Graveley et al. 2011; Lu et al. 2010). The new Araport11 annotation consists of 37,592 genes (27,688 protein-coding, 5,051 noncoding, 952 pseudogenic, and 3901 transposable element-related loci) and 58,149 transcripts. The annotated loci span 67.7 Mb (56.6% of the genome sequence), an increase from 61.2 Mb (51.2%) in TAIR10.

The updated annotation revealed a sophisticated landscape of the transcriptional structures of genes, in terms of their splicing patterns and UTRs. The number of protein-coding genes expressing multiple splice isoforms increased from 21% in TAIR10 to 39% in Araport11. This figure is lower than a previous estimate of 54%, reported as 61% of intron-containing genes are alternatively spliced (Marquez et al. 2012). Therefore, we analyzed the same normalized cDNA libraries made from mixed flowers and seedlings (Marquez et al. 2012) using the Araport11 pipeline and found ~16% of genes have more the one splice isoform. This indicated that the criteria used in the two studies played a major role in the discrepancy. Nevertheless, we believe the annotation of splice isoforms is not saturated yet for two reasons. First, we only interrogated datasets generated from wild-type plants under regular growth environment. Thus splicing events that occur only under stressed conditions and/or mutants may not be present and annotated. Second, PASA explicitly requires canonical splice sites and eight perfect matches on each side of a splice site to minimize incorrect transcript structures. Enforcing these filters is likely to have missed some splice variants with non-canonical splice sites.

Analysis of the exitrons identified in this work overlapped 209 of those previously reported (Marquez et al. 2012). We investigated the 793 exitrons exclusively detected by Marquez *et al.* and found that in those loci, the exitron-retained transcripts, but not the exitron-spliced counterparts, were annotated in Araport11. This indicated that the exitron-retained transcripts in those loci were the major isoforms. Taken together with our results showing that the ribosome occupancies of exitrons were comparable to those of coding exons (Fig. 4E,F), splicing out of exitrons may be considered as a means of exon skipping behavior with post-transcriptional regulatory functions.

Approximately one-third of the splicing activities occur in the UTRs and do not affect protein sequence. For example, the phytochrome-interaction factor 3 (*PIF3*) encodes six transcript variants with identical coding sequence; three of these mRNA isoforms were specifically present in dark-grown seedling samples. The complex UTR landscape we captured in Araport11 is in agreement with the heterogeneity of transcription start sites and polyadenylation sites (Sherstnev et al. 2012; Morton et al. 2014). In addition, we noted that many genes overlap at their boundaries, a feature also observed in other eukaryotic species (Wang et al. 2009). One challenge we encountered was that the short read lengths (<=100 bp) and non-strandedness of the 113 RNA-seq datasets make a completely automatic pipeline inadequate to provide unambiguous gene boundaries for overlapping genes. We found manual curation with consideration of additional datasets a necessary and worthwhile effort to avoid invasive UTR annotations. Likewise, increasing effort has also been placed on manual curation over the past decade to maintain the most trusted metazoan genomes (Schurch et al. 2014). Most of the 508 novel transcribed regions we identified were not reported by comparative sequence analysis. This reinforces the importance of using transcriptome profiling to complement comparative analyses for genome annotation (Haudry et al. 2013), similar to the phenomenon observed in fly (Graveley et al. 2011). Interestingly, many of the NTRs that overlap with conserved noncoding sequences are also occupied by ribosomes in our analysis. Given that ribosome occupancy alone is insufficient to classify transcripts as coding or noncoding (Guttman et al. 2013), the next challenge is to uncover whether these NTRs encode small peptides as found in Arabidopsis and other species (Lauressergues et al. 2015; Quinn and Chang 2015; Cohen 2014).

Despite the substantial increase in structural diversity and expression profiling presented in Araport11, the annotation of *Arabidopsis thaliana* should be an ongoing project. Additional samples from developmentally specific stages (i.e. embryonic and senescent tissues) and environmental stimuli and advancement in long read sequencing technology will most certainly result in the discovery of novel features. Furthermore, the small RNA annotation demonstrates that the integration of published datasets and analytic tools could provide a comprehensive and uniform annotation for the research community. Finally, advances in bioinformatics in the analysis of the ribosome profiling data will provide valuable new insights into the translational activities and classification of both novel and existing transcripts.

## METHODS

### Datasets

We assessed over 400 RNA-seq datasets available as of September 2014 from the NCBI SRA. Among these, 113 Illumina-based datasets generated from wild-type Col-0 ecotype untreated or mock-treated samples were used in the annotation pipeline. A complete list of SRA accessions with detailed sample descriptions is available at Araport (www.araport.org/rna-seq-read-datasets-used-araport11). We describe the processing steps in detail in the Supplemental Methods.

### Tissue-specific transcriptomes

To instantiate tissue-specific transcript isoforms and avoid possible chimeras between tissues, each tissue bin was assembled independently using Trinity (release 20140413) (Grabherr et al. 2011) using both *de novo* and genome-guided modes. For the latter, only uniquely mapped reads were used by Trinity to guide the *de novo* assembly of overlapping read clusters at each locus. Both procedures enforced two cutoff values: a minimum contig length of 183 bp, as 95% of Arabidopsis transcripts are greater than or equal to this length, and a maximum intron length of 2000 bp which is the 99th percentile for intron sizes in TAIR10 gene models. The separate *de novo* and genome-guided results were collapsed, mapped back to the reference genome and merged into compatible transcript clusters by PASA (Haas et al. 2003) using its alignment assembly module. These transcript clusters were sub-assembled into compatible transcript structures, which adhered to the following criteria: maximum intron length of 2000 bp, ≥ 90% of transcript aligned at ≥ 95% identity, both sides of the splice boundary are supported by at least 8 bp and the splice sites are canonical. We describe refining approaches in the Supplemental Method “Structural annotation refinement”.

### Novel transcribed regions

Overlapping novel transcripts localized within Araport11 intergenic regions were clustered into novel transcribed regions. We used featureCounts (v1.5.0-p1) (Liao et al. 2014) to calculate the number of reads aligned to the novel transcribed regions in the 113 RNA-seq datasets. We only retained the regions that either contained more than one read count per million mapped reads in any given tissue or encode intron-containing transcript(s). Overall, we proposed 508 novel transcribed regions in Araport11 and their transcript models.

### Annotation edit distance

We used MAKER (2.31.8) (Holt and Yandell 2011) under default settings to compute AED for each transcript of protein-coding genes, noncoding genes, and pseudogenes. We supplemented the following evidence: 1) PASA-assembled transcripts generated in this study, 2) the EST, full-length cDNAs and RNA-seq data including non-control and additional ecotype samples generated by MAKER-P (Campbell et al. 2014) and 3) UniRef90 protein sequences for ten plant species (see “Novel transcribed regions” in Methods) aligned to TAIR10 genome using the protein2genome option of MAKER.

### Small RNAs

We used Cutadapt (Martin et al. 2010) to trim the reads from 9 studies (Supplemental Table S2) and identified *de novo* small RNA clusters (miRNA search disabled,-nohp) using ShortStack (v3.3) (Axtell 2013). The reads from replicate samples were coupled in a single ShortStack analysis, and only the *DCL*-derived loci called out by ShortStack were retained for the remaining analyses. We used BEDtools (Quinlan and Hall 2010) to merge overlapping clusters from different studies and adopted the consensus regions as the small RNA reference. ShortStack computes the dominant 20- to 24-nt small RNA size for each locus where mixtures of different sized RNA are possibly generated (Coruh et al. 2015). During the merging step, we used the predominant RNA size within each consensus cluster, and designated “uncharacterized” (Fig. 2A) if a dominant size was not available. We computed the small RNA abundance using ShortStack count mode under default settings for wild-type, *nrpd (pol-iv)*, *nrpe (pol-v)* libraries (Fig. 2B) made from stage 1-12 flowers (Law et al. 2013). The circular visualization (Fig. 2C) was generated using the circlize package (Gu et al. 2014). We used Bedtools (Quinlan and Hall 2010) to compare the genomic regions of the small RNA clusters with the P4RNA loci (Li et al. 2015; Zhai et al. 2015).

### Expression profiles

Gene expression values were computed on 113 RNA-seq datasets at the gene and transcript levels using Salmon (v0.6.0) quasi-mapping-based mode and the variational Bayesian EM algorithm (Patro et al. 2015). The expression data are available at Araport and also via ThaleMine 1.9.0. Addition information about expression analyses is available in the Supplemental Methods.

### Alternative splicing events

We calculated alternative splicing events on Araport11, TAIR10 and human (Ensembl release 83) annotation using the generateEvents operation of SUPPA (v1.2b) (Alamancos et al. 2015). To calculate the PSI value for each splicing events, we ran the psiPerEvent operation of SUPPA using Salmon-generated expression file as input. We compared the PSI value distributions of retained intron events between wild-type and mutant samples using Wilcoxon rank sum test, removing events with missing PSI values.

## DATA ACCESS

Araport11 has been submitted to NCBI GenBank under the accession numbers CP002684 (Chr1), CP002685 (Chr2), CP002686 (Chr3), CP002687 (Chr4), and CP002688 (Chr5). The corresponding NCBI BioProject and BioSample numbers encapsulating the above GenBank accessions are PRJNA10719 and SAMN03081427 respectively. The data can also be accessed via Araport and a list of links to each dataset is available in Supplemental Data Set S4.

## ACKNOWLEDGMENTS

We would like to thank Michael Axtell, Eduardo Eyras, and Carson Holt for helpful discussions on software utilization (ShortStack, SUPPA, and MAKER-P), Kim Pruitt and Ho-Ming Chen for constructive comments, and Jason Miller and Sergio Contrino for critical reading of the manuscript. This work was funded by the National Science Foundation (DBI-1262414).

